# An integrative web-based software tool for multi-dimensional pathology whole-slide image analytics

**DOI:** 10.1101/2022.06.08.495366

**Authors:** Alice Shen, Fusheng Wang, Saptarshi Paul, Divya Bhuvanapalli, Jacob Alayof, Alton B. Farris, George Teodoro, Daniel J. Brat, Jun Kong

## Abstract

In the era of precision medicine, human tumor atlas oriented studies have been significantly facilitated by high-resolution, multi-modal tissue based microscopic pathology image analytics. To better support such tissue-based investigations, we develop Digital Pathology Laboratory (DPLab), a publicly available web-based platform, to assist biomedical research groups, non-technical end users, and clinicians for pathology Whole-Slide Image (WSI) visualization, annotation, analysis, and sharing via web browsers. A major advance of this work is the easy-to-follow methods to reconstruct three-dimension (3D) tissue image volumes by registering two-dimension (2D) whole-slide pathology images of serial tissue sections stained by hematoxylin and eosin (H&E), and immunohistochemistry (IHC). The integration of these serial slides stained by different methods provides cellular phenotype and pathophysiologic states in the context of a 3D tissue micro-environment. DPLab is hosted on a publicly accessible server and connected to a backend computational cluster for intensive image analysis computations, with results visualized, downloaded, and shared via a web interface. Equipped with an analysis toolbox of numerous image processing algorithms, DPLab supports continued integration of community-contributed algorithms and presents an effective solution to improve the accessibility and dissemination of image analysis algorithms by research communities. It represents the first step in making next generation tissue investigation tools widely available to the research community, enabling and facilitating discovery of clinically relevant disease mechanisms in a digital 3D tissue space.

## INTRODUCTION

Significant advances in high-throughput histologic slide scanning technologies have enabled imaging and analyses at cellular and subcellular levels and have allowed high-resolution characterizations of cell and matrix structure profiles, tissue microenvironments and biomarker distributions [1-6]. However, the extreme size of the resulting giga-pixel images presents tremendous performance and analytic challenges. Traditional manual analyses are not practical due to the overwhelming image scale and substantial inter-and intra-reader variability. By contrast, automated computerized image analysis algorithms have emerged to accurately assess clinically relevant pathologic features in high-resolution pathology WSIs.

Despite methodologic improvement, the potential for capturing and analyzing the rich information embedded in WSI data has not been fully realized. A major limiting factor is that traditional computer software and file sharing methods are inadequate for sharing, visualizing and analyzing giga-pixel images. While numerous offline image processing tools have been developed, such as Fiji [7], Icy [8], CellProfiler [9], among others, they are primarily developed for reasonably sized or cropped 2D sub-regions of WSIs. End users of these tools rely on local computational and storage resources, which are often insufficient. Thus, such lack of connection to large-scale computational facilities necessary for high-throughput image processing limits the usage of such tools for pathology WSI analysis.

In addition, almost all WSI visualization and analysis tools are constrained by the 2D image space [10-15]. As 2D projected spatial profiles of the 3D pathologic features depend heavily on planes selected during tissue processing and slide preparation, the resulting morphological and spatial characterizations are subject to significant information loss, bias, and even distortion. Thus, the ability to create, visualize and analyze high-resolution 3D tissue volumes would represent a significant step forward in tissue-based biomedical investigations.

A further step towards a more comprehensive research strategy for tissue microenvironment characterizations would integrate histopathologic images with spatially mapped images of pathophysiological biomarkers in a single 3D tissue space. This combination would provide multi-modal and multi-scale information on cellular phenotype, signaling networks, and physiological state with well-preserved spatial relations. Such technology would be highly beneficial for better understanding cellular interactions that drive tissue development, homeostasis, and disease progression.

To enable such enhanced tissue-based biomedical investigations, we develop an integrative web-based platform to support multi-dimensional pathology WSI analytics. Web applications are widely available and accessible from web browsers without installations or configurations. Web-based frameworks facilitate the storage and portability of large amounts of data, as uploaded files are stored in a central cloud server, reducing the storage burden of individual users and the need for sustained local storage of large datasets. Large data sets can be made accessible to others without copying or transferring datasets, and the obstacle of sharing large files with collaborators becomes less relevant. Importantly, web applications do not require powerful local computational resources. A remote web application architecture allowing users to access a shared cluster of computational facilities addresses numerous computational challenges in pathology WSI analysis. The developed web application also enables the capability to create 3D tissue volumes from 2D serial tissue slides, including the incorporation of interdigitated biomarker sections [16-19]. Our online platform represents a substantial step forward for realizing the full potential of high-resolution integrative 3D tissue analytics by the research community [20–23].

## RESULTS

### Application overview

We have developed Digital Pathology Laboratory (DPLab) platform (https://dplab.gsu.edu/), the first publicly available online “lab-in-a-browser” informatics platform, to facilitate the management, annotation, sharing, visualization, integration, and analysis of WSIs in the 2D and 3D virtual tissue space. All functionalities of DPLab, including data collection, visualization, annotation, image processing, and sharing can be accessed via a point-and-click web interface, allowing users to conduct and collaborate on image analysis experiments from a web browser.

The DPLab removes technological barriers that hinder the use of high-resolution WSIs and advanced analysis algorithms by supporting all pipeline steps via a point-and-click web interface backed by a remote computational cluster. The platform supplies computational resources necessary to run analyses on giga-pixel WSIs, as analyses are performed on a remote computational server and pose no computational load to the client machine. Using the web interface, users can configure, queue and run an array of image analysis algorithms without writing additional configurations or scripts. Upon completion, users can visualize analysis result sets, screen results with interactive corrections, and download result data for further correlative and statistical analyses. The platform fully supports numerous image modalities widely used in tissue-based research, including pathology WSIs in SVS and NDPI formats, and radiology MRI image volumes. Other commonly used image formats, such as JPG, TIF, and PNG, are supported. The platform is currently deployed with a suite of publicly available algorithms, including cell seed detection (Deep Learning-based and Non-Deep Learning methods), nuclei segmentation, IHC positive pixel counting, color normalization, fibrosis quantification (Deep Learning-based and Non-Deep Learning methods), steatosis segmentation (Deep Learning-based and Non-Deep Learning methods), DAPI signal based nuclei segmentation from Imaging Mass Cytometry data (Deep Learning), and dynamic serial WSI registration. The library of algorithms is extensible and includes a built-in API to facilitate future contributions from the research and scientific community. Functionality currently available on the DPLab platform is listed in Table 1 in comparison to other available state-of-the-art pathology image visualization and management software [24-27]. All key functions provided by DPLab web application are discussed in depth in the Results section and demonstrated in videos posted on a web page included in Supplementary Materials.

**Table 1.** Comparison of functionality in DPLab with that from other state-of-the-art software options.

| **Functionality** | **Aperio**<br>**Webscope<sup>41</sup>** | **Hamamatsu**<br>**NDP<sup>42</sup>** | **Omero<sup>43</sup>** | **QuPath<sup>44</sup>** | **Digital Pathology** |
| --- | --- | --- | --- | --- | --- |
| In-Browser | Yes | No | Yes | No | Yes |
| Cloud Storage<br>and Hosting | No | No | No | No | Interfaced to cloud |
| Image Upload | Yes | Yes | Yes | Yes | Yes |
| Support for 2D<br>Image Visualization | Yes | Yes | Yes | Yes | Yes, and 3D |
| Support for Serial<br>Pathology WSIs | No | No | No | No | Yes |
| Support for<br>Integration of<br>Images of Different<br>Modalities | No | No | No | No | Yes |
| Image Annotation | No | No | Yes | Yes | Yes |
| Image Analysis | No | No | Yes | Yes | Yes |
| Result Visualization<br>and Download | No | No | Yes | Yes | Yes |
| Interactive Result<br>Filtering and<br>Selection | No | No | No | No | Yes |

### User account creation and verification process

The DPLab web application requires a user to create an account with a valid email. After an account request is submitted by a user, a confirmation email is sent to the user. In the meanwhile, a separate approval email is sent to the DPLab administrators to prevent automated malicious threats for enhanced data security. After authorization, an approval email is sent to the user. Requests to retrieve or reset passwords can be initiated at the login page. This complete process is demonstrated by Supplemental Movie S1 on user account creation. With web security concerns, user account requests need to be approved by DPLab administrators before users can use the DPLab platform.

### WSI upload and 2D WSI visualization

The DPLab platform supports the typical end-to-end pipelines and workflows to perform image analysis on either pre-or user-uploaded image sets, beginning with image upload and visualization. Users can upload de-identified images privately to their accounts or contribute images to the searchable public image database. After upload, users can use the in-browser visualization interface to conveniently view and navigate across their data using natural mouse gestures and interface buttons. For reviewing WSIs, users can click or scroll to zoom and drag to pan across such high-resolution images at various magnifications. This interface design mimics the way pathologists review physical slides under a microscope. The interface enables visualization of a WSI at the maximum resolution without significant computer memory consumption by using a Deep Zoom Image (DZI) pyramidal tiling schema to serve and display tiles of images only within the current field of view at the requested resolution [28, 29].

### WSI management

Users can create multiple projects to manage WSI data for different studies. Each project can be assigned with multiple project properties for project description. A demonstration is given by Supplemental Movie S2 on project management. Additionally, DPLab allows users to search and explore images at the project level based on various attributes, such as the project title, disease site (e.g. brain, liver, etc.), imaging modality (e.g. CT, MRI, etc.) and image data dimensionality (e.g. 2D, 3D, etc.). Similarly, the platform allows users to search images within a project based on image title, image type (e.g. 2D, 3D, etc.), annotation class, and annotation label. The uploaded images can be moved across projects if the user decides to restructure projects. Users can also delete images, projects, annotations and analysis results. Additionally, DPLab supports easy sharing of projects or analysis results among users. A demonstration on image management is given by Supplemental Movie S3.

### Annotations on WSIs

Supported by the visualization interface, users can make flexible annotations on each image. Specifically, users can either annotate a single region of interest (ROI) for image analysis or multiple ROIs for algorithm training. Multiple annotation classes can be created to better describe ROIs. Each annotation class can include multiple ROIs of different types, including rectangles, circles, points, and free hand contours. A detailed annotation process is demonstrated by Supplemental Movie S4 on ROI annotations. The resulting ROI annotations can be downloaded and shared online with other research and clinical collaborators who have accounts on the DPLab platform. The annotations can be downloaded as either a JavaScript Object Notation (JSON) or an Extensible Markup Language (XML) document. Pre-generated annotations from other sources that conform to a predefined JSON format can also be uploaded to the DPLab database. To support better integration of other common annotation formats, we have also implemented compatibility with Aperio ImageScope compliant XML annotation files. This compatibility allows users to re-use pre-collected annotations made by Aperio ImageScope, one of the most popular annotation tools widely used in the tissue image-based research community [30]. It also supports to produce Aperio ImageScope XML annotation files from DPLab platform directly. In this way, annotations can be shared seamlessly between DPLab platform and Aperio ImageScope in bilateral directions. The compatibility of annotation files to Aperio ImageScope XML format is demonstrated by Supplemental Movie S5 on annotation files.

### Digital tissue 3D reconstruction and integration by serial WSI registration

As pathologic diseases occur in the 3D space, structural, and spatial information derived from 2D imaging data is significantly diminished and even distorted from the 3D space. However, 3D tissue volume reconstruction with serial histopathology WSIs is technically challenging due to the giga-pixel image scale, the presence of millions of micro-anatomical objects, and large tissue variations. We have deployed to DPLab our dynamic multi-resolution registration method that allows users to efficiently scrutinize volumes of interest on-demand over the entire WSI domain at the full cellular resolution in a 3D tissue space without expensive computation [31].

Two result sets with serial WSIs of Glioblastoma and Liver tissue sections in H&E and Masson’s trichrome stain are presented in Figure 1. For each WSI, we first extract its low-resolution representation. With low-resolution images, global tissue structures are used for macro-level registration with global landmark-based transformations. DPLab supports user-provided landmarks both from uploaded files and DPLab annotation interface. Additionally, DPLab can detect landmarks by Speeded Up Robust Feature (SURF) detector and match landmark pairs with local descriptors. Due to image artifacts (e.g. tissue folding, out of focus, tissue contamination by marker pens, among others), we overlay resulting identified point pairs with low-resolution image pairs and allow users to correct them before transformation estimation. Each landmark pairs are labeled by unique numbers. Users can add new landmarks, move or delete existing point pairs by DPLab provided web interface. As all these analyses use low-resolution image representations, image data can be readily fit to machine memory for efficient processing. Next, we estimate an optimal similarity transformation with purified landmark pairs at low-resolution and map such transformation to the highest image resolution for dynamic registration of informative regions at the full image resolution. This multi-resolution based solution addresses the computer memory overflow problem. For serial image registration at a high image resolution, we implement an iterative transformation propagation method to align non-reference images to a common reference image [31]. The resulting registered images are combined as an image volume for web-based review. We visualize the high-resolution serial WSI stacks before registration, extracted low-resolution serial WSI stacks before registration, resulting low-resolution WSI stacks after the low image resolution registration, and high-resolution serial image stacks after the high image resolution registration in a 3D virtual tissue space in Figure 1(A-D), respectively. A step-by-step demonstration of the whole registration process is given by Supplemental Movie S6 on WSI registration analysis.

**Figure 1.**
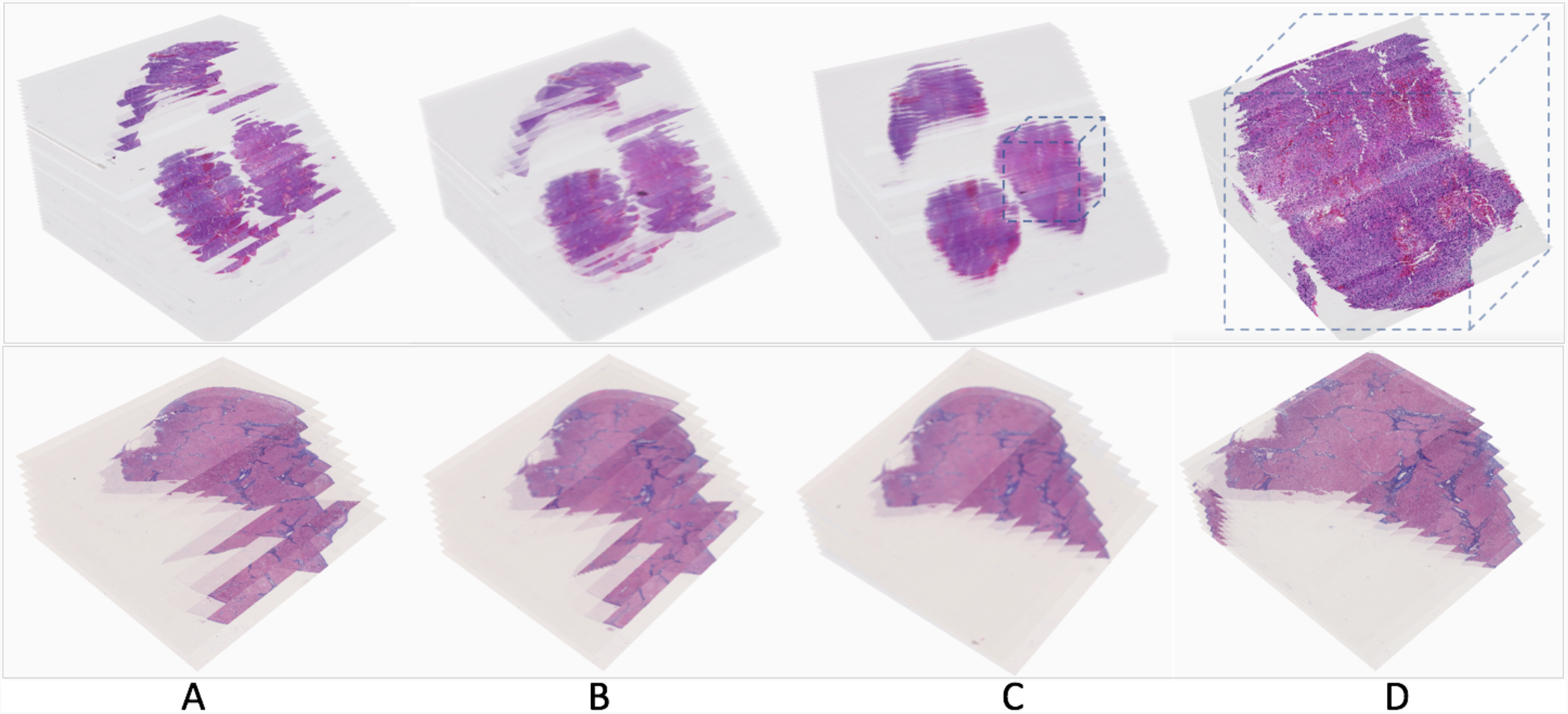
Serial pathology image registration results. Two result sets with serial WSIs of (Top) Glioblastoma and (Bottom) Liver tissue sections in H&E and Masson’s trichrome stain are demonstrated. (A) High-resolution serial WSI stacks before registration, (B) extracted low-resolution serial WSI stacks before registration, (C) resulting low-resolution WSI stacks after the low image resolution registration, and (D) high-resolution serial image stacks after the high image resolution registration, are visualized in a 3D virtual tissue space.

### Tissue visualization in a 3D virtual tissue space

DPLab fully supports diverse image modalities and provides interfaces for users to visualize and navigate through image stacks in 3D space. Users can either visualize each slide of a volume as a 2D image or choose to visualize the entire volume in a 3D viewer. To our best knowledge, the ability to view a serial image stack as a 3D image volume is unique to DPLab and is not currently available in any other web-based digital pathology tool. Powered by WebGL and ThreeJS JavaScript libraries, the 3D DPLab visualization tool combines each slide of an image stack and constructs serial pathology slides into a 3D volume for in-browser visualization, allowing users to navigate through the image volume by natural mouse gestures for rotation, panning, and zooming [32, 33]. The interface also offers an inbuilt tool allowing users to clip, or “slice” into 3D image volumes for better insight on spatial characteristics of their data in 3D space. The clipping plane is fully adjustable and can run along vertical (y), horizontal (x), depth-wise (z) or diagonal axes. Figure 2 illustrates the 3D tissue volume visualization tool with registered serial tissue regions. Users can adjust the opacity of each image slide for enhanced visualization of tissue histological and spatial features in the image volume, as well as adjust slide spacing for more extended or condensed visualizations. The interface also provides options to display image boundaries, axes and planes for ease of visualization. Tools are available to display an adjustable marker boundary box to highlight and outline ROIs within 3D space. A detailed use of this 3D visualization tool is demonstrated by Supplemental Movie S7 on 3D visualization.

**Figure 2.**
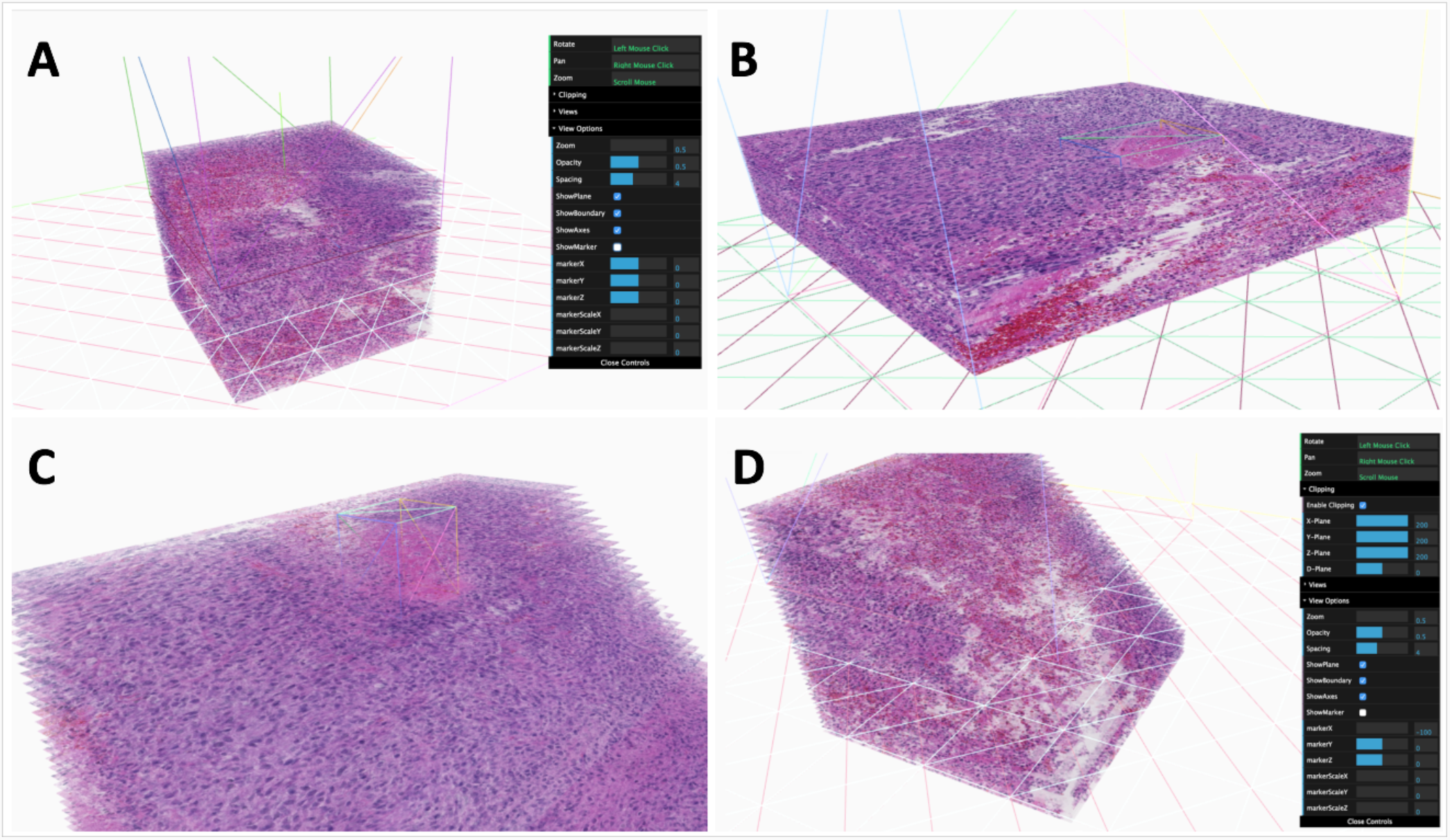
3D Visualization and slicing interface. Users can visualize imaging data in 3D volumes directly from a browser. The 3D visualization interface allows for a variety of mouse gestures and adjustable settings to pan, zoom and slice/cut into image volumes. Options for opacity adjustment, slice spacing/thickness, as well as 3D guidelines and markers are also available for use. (A) Image volume at a thickness of 3 units, with gridlines and boundaries visible; (B) Image volume at a thickness of 1 unit, with gridlines and boundaries visible; (C) Image volume with an adjustable 3D marker placed around a feature of the image stack; (D) Image volume with diagonal clipping, revealing features displayed by cross-sectional views within the volume.

DPLab supports 3D visualization of 3D image volumes in numerous image formats, including serial histopathology slides stained with H&E, serial histopathology slides stained with IHC, serial slides alternatively stained by H&E and IHC, as well as radiology image volumes (both MRI and CT modality). We illustrate sample visualizations of these different image modalities in Figure 3.

**Figure 3.**
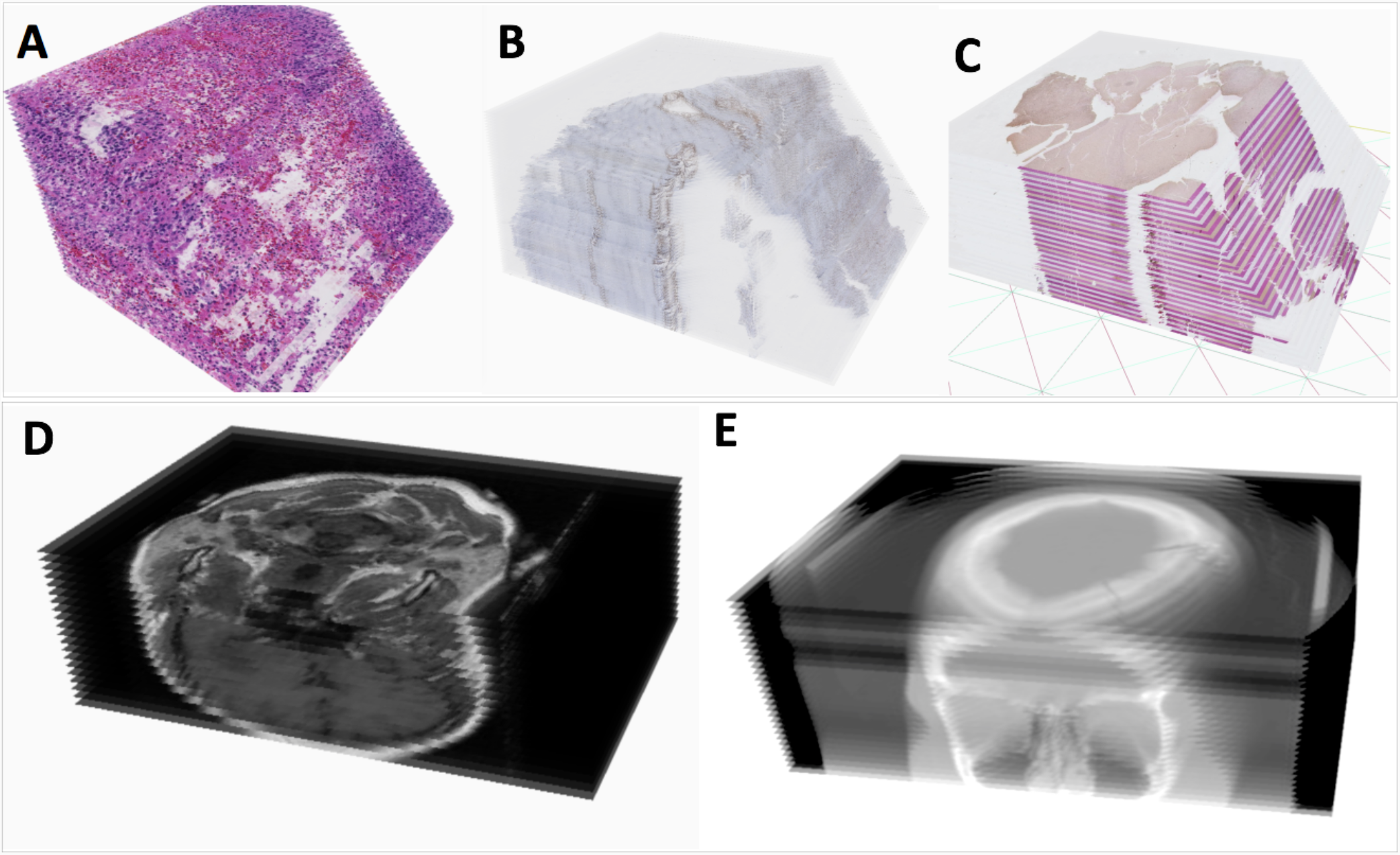
3D Visualization support for multiple image modalities. (A) Registered serial H&E WSIs; (B) Registered serial IHC WSIs; (C) Registered serial WSIs stained by H&E and IHC biomarker alternatively; (D) Digital Imaging and Communications in Medicine (DICOM) radiology serial image volume of a patient with a brain neoplasia; and (E) DICOM CT serial image volume of a patient, with sinus cavities visualized.

### Algorithm invocation and library expansion

Through a simple web form, users can select an algorithm from a library of contributed algorithms and configure parameters (either default or customized parameter values) to run analyses on WSIs or ROIs drawn within a selected image. We demonstrate the general procedure to invoke and submit analysis jobs by Supplemental Movie S8. After the analysis request is queued, a remote computational server performs tile-wise analyses of the selected algorithm on the WSI or ROI and stores the results in a format suitable to the output modality of the algorithm. A 3D image processing algorithm yields results stored as a series of 3D coordinates, which are next visualized as 3D voxels or meshes over various slides, while 2D results are visualized as points, lines or polygons. Algorithm results are stored in the database and can be visualized directly within the browser as polygons or masks overlaid on the original image. Results can also be shared with collaborator accounts or downloaded in a raw data format for extended offline use.

A suite of algorithms deployed to the DPLab platform is available for public use, as listed in Table 2. Examples of the result visualizations of 2D algorithms for nuclei segmentation, IHC positive pixel counting, liver fibrosis quantification, cell seed detection, steatosis segmentation, and nuclei segmentation from multiplexed Hyperion data are shown in Figure 4, respectively. The opacity of object contours in segmentation results can be changed by users to facilitate result review. For IHC positive pixel counting, the strong, medium, and low positive pixel percentages are provided. Additionally, users can choose to display only strong, medium, or low positive pixels. For the liver fibrosis quantitation, the numerical fibrosis percentage is reported in addition to the visual processing results.

**Table 2.** A list of image analysis algorithms available on the DPLab platform.

| Title | Input Data Type | Output Data Type | Language | Description |
| --- | --- | --- | --- | --- |
| Cell Seed Detection (Conventional methods) | 2D Images | Points | MATLAB | Detection of cells with information from spatial connectivity, edge map, and a voting mechanism with a local shape-based intensity analysis |
| Cell Seed Detection (Deep Learning) | 2D Images | Points | Python, C, C++ | Detection of cells using Convolutional Neural Network (CNN) with interactive learning |
| Cell Segmentation (Hysteresis Thresholding) | 2D Images | Polygonal ROIs | MATLAB | Segmentation and division of cells with Hysteresis analysis and watershed method |
| IHC Positive Pixel Counting | 2D Images | Polygonal ROIs, Percentages | MATLAB | Identification of pixels with strong, intermediate, and weak stain intensity |
| Steatosis Segmentation and Segregation (Conventional methods) | 2D Images | Polygonal ROIs | MATLAB | Segmentation and segregation of liver steatosis for quantification |
| Steatosis Segmentation and Segregation (Deep Learning based) | 2D Images | Polygonal ROIs | Python | Segmentation and segregation of liver steatosis for quantification |
| Dynamic Serial Slide Registration | 2D Serial WSIs | 3D Image Volume | MATLAB | Dynamic registration of user-specified regions from whole-slide serial pathology image sequence with multi-stage and multi-resolution mapping and aggregation methods |
| DAPI Nuclei Segmentation | 3D Multi-Channel Image | 2D Image of Components | Python | DAPI signal based nuclei segmentation by deep learning with Imaging Mass Cytometry data |
| Fibrosis (Conventional methods) | 2D Images | Polygonal ROIs | MATLAB | Identifications of liver fibrosis regions with contours |
| Fibrosis (Deep Learning based) | 2D Images | Polygonal ROIs | Python | Identifications of liver fibrosis regions with contours |
| Color Deconvolution | 2D Images | 2D Images | Python | Color deconvolution/unmixing for HE, IHC or other customized stain images |
| Color normalization | 2D Images | 2D Images | Python | Normalization of color with HE, IHC, or other customized stain images |
| White Area Segmentation | 2D Images | 2D Images | Python | Segmentation of white areas within tissue images |
| Extract region of interest from WSIs | 2D Images | 2D Images | Python | Annotation, extraction, and download of regions of interest from a WSI |

**Figure 4.**
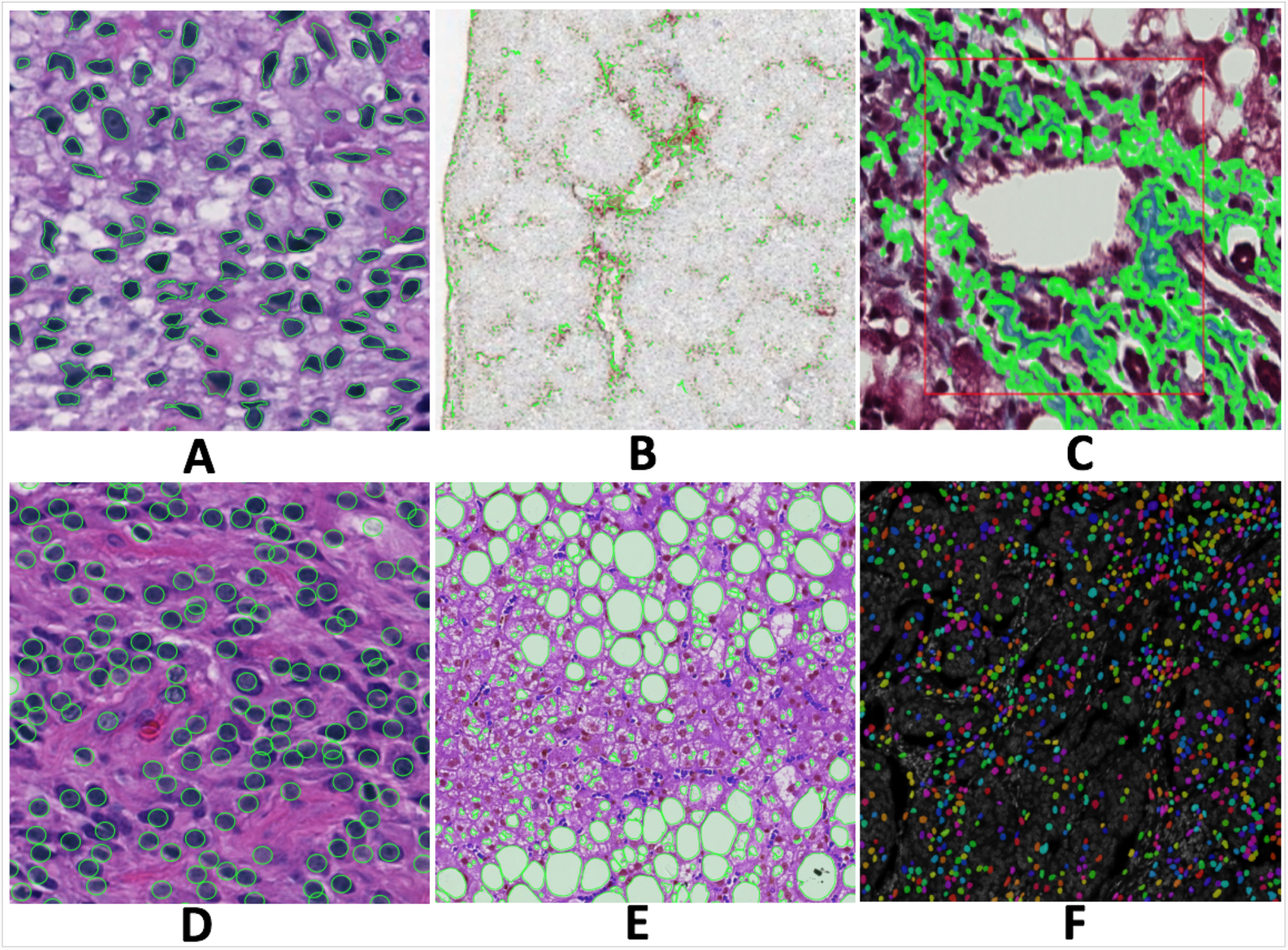
Typical 2D image analysis result visualizations from deployed 2D algorithms. Algorithms return green contours and masks in colors to represent detection and segmentation results. (A) Nuclei segmentation; (B) IHC positive pixel counting; (C) Fibrosis quantification; (D) Cell seed detection; (E) Steatosis segmentation; and (F) Nuclei segmentation with multiplexed Hyperion biomarker data.

The algorithm library is designed to be continuously expanded by community contributions. New algorithms can be uploaded to the platform and made available for public use via an open API. Supported by an easy-to-access and code-free interface, the open algorithm library on DPLab is designed to help distribute and validate new algorithms and promote their usage by both researchers and clinicians in collaborative contexts. Algorithms are contributed to DPLab by submissions of their code, any documentation or accompanying literature for review by the DPLab platform development team. If the algorithm code is functional and is correctly formatted according to API standards, it will be incorporated into the DPLab library and interface and made available for public use. The algorithm library API currently supports a variety of commonly used languages, including Java, MATLAB, Python, Julia, C, and C++. However, the languages supported by DPLab are adaptable and can be extended as the library grows. We demonstrate how to integrate and manage new algorithms to expand the analysis library on DPLab by Supplemental Movie S9 on algorithm deployment.

### Iterative algorithm training with refined results

In addition to running analysis algorithms on 2D images and 3D image volumes, DPLab allows visualization of results directly within the browser without the need for any additional scripting, pipeline coding, or computational infrastructure access or configuration. During in-browser visualization, users can click on contours or points generated from results to include or exclude subsets of results from export. This is particular useful for supervised machine learning algorithms that can be readily refined with the result visualization and selection interface. With this interface, users can conveniently remove manually false positive or incorrect results by clicking on the undesired contours. Excluded contours will turn red to visually distinguish them from the included green contours. Refined result datasets can then be exported and used to retrain machine learning algorithms, after which the retrained algorithm can be re-uploaded to DPLab to generate new result datasets. This process can be repeated multiple times to iteratively improve the quality of training datasets, thus improving the accuracy and selectivity of machine learning algorithms. We demonstrate this interactive process for training liver steatosis detection algorithm in Supplemental Movie S10. Using this functionality, supervised machine learning algorithms, such as Deep Learning-based methods, can be efficiently re-trained and refined in a user-mediated, iterative manner.

## DISCUSSION

Currently available digital pathology tools and platforms still present challenges for users to reliably use such tools in real-use applications, as they provide limited critical functionality for state-of-the-art data upload, analysis, and visualization, and have high technical set up costs that are prohibitive. DPLab is developed as a solution to address these limitations. It provides comprehensive and accessible tools to facilitate end-to-end pathology image analysis pipelines for a variety of potential use cases in research and clinical settings.

Recently emerging 3D digital pathology is enabled by the spatial registration of digital images of serial 2D tissue sections. 3D tissue assessment, especially with the incorporation of biomarkers, has substantial potential for understanding development, physiology and disease, since it allows visual reviews of a tissue in its entirety rather than selected, and potentially misrepresentative 2D representations. Such analyses present strong translational values and are promising to support clinical practice in pathology for specific disease states like breast cancer and prostate cancer. Certainly, in the setting of research and education, these 3D applications can better inform pathologists and investigators on tissue features and in-situ biomarkers. The DPLab web platform includes novel image registration methods that dynamically align multi-resolution giga-pixel WSIs of serial tissue sections, a key step for transforming information from extremely large serial images to 3D spatial objects. This method also enables integration of adjacent H&E stained images providing micro-environmental context with spatially mapped IHC images that capture cellular phenotype and pathophysiological properties.

Given the time-consuming nature of pathology image annotation, DPLab can also serve as an efficient way to facilitate the composition of high-quality training datasets for supervised machine learning algorithms, e.g., the deep Learning based pathology image analysis. Researchers can upload their image data and perform analyses using a supervised machine learning-based algorithm currently available in DPLab built-in algorithm suite. After analyses are completed, researchers can use the DPLab result visualization interface to interactively select and download high-quality subsets of the generated results, which can then be used to re-train and improve the accuracy of subsequent supervised machine learning algorithms. In this way, DPLab substantially reduces time cost to establish high-quality, large-scale training datasets, a common problem in deep Learning-based methods, and also initiates a vision-guided interactive and iterative training mechanism applicable to a broad set of machine learning algorithms for enhanced performance.

Considering the wide variety of bio-image data, we are actively expanding support on the platform for more data formats and modalities. As the DPLab platform evolves, integrated support for more modalities of multi-dimensional data, including temporal data volumes and fluorescence microscopy image data will be incorporated. Support for genomic data uploading is also under active development, with the goal of developing functionalities for more integrative analysis with clinical, genomic, and phenotypic results derived from image analysis. Future work on built-in statistical methods will allow for more powerful data integration and analysis. Histological features derived from analyses, such as 2D/3D morphology and spatial organization features, can be represented with high order statistical descriptors, which can then be used for studies on outcome prediction [6, 34-39].

The DPLab application is supported on privately hosted servers. As demand grows, we foresee a migration onto cloud-based infrastructure, such as Amazon Web Services (AWS), Google Cloud or Microsoft Azure [40–42], to support higher availability and resource scalability. With this being the long-term goal, the design patterns and architecture utilized in the creation of this application are intended from the outset to support high scalability and cloud-based hosting. Due to the Javascript libraries used by DPLab, the full functionality of the application can only be supported by the Google Chrome browser. For a better user interface experience, we plan to move the view from MVC framework to Angular or React [51–52], resulting in a higher application scalability.

To encourage researchers and clinicians to use DPLab, we plan to support an increasingly diverse range of algorithms written in commonly utilized programming languages and solicit contributions from the image analysis algorithm community, which will include detailed and easy-to-follow documentations on how to contribute and share user-contributed algorithms to the DPLab library via the pre-specified API standards. We also have plans to develop iOS and Android applications for mobile and tablet devices for an extended and more convenient user experience, especially for clinicians in clinical environments.

## METHODS

The web application is primarily built in Ruby on Rails (RoR) [43], a widely used and well-proven web framework with industry-standard design patterns (Model-View-Controller) that has been used to create dynamic, robust, and scalable web applications such as Groupon [44], Github [45], Airbnb [46], and many other high-load, high-performance web applications used by millions of people worldwide. The primary data tables containing all application related data (e.g., users, images and other model objects) are stored within a PostgreSQL database. The primary database system, PostgreSQL, is chosen for its high performance in applications with large amounts of data (e.g., scalability, synchronous replication, rich indexing), flexibility, and extensible data types. These signature properties allow our application to evolve, as new forms of image modalities and algorithms are incorporated [47]. The various functionalities made available by the front end of the web application, such as 2D and 3D image visualization and annotation drawing utilize numerous JavaScript libraries and APIs, including OpenSeadragon [28], OpenSlide [48], ThreeJS [32], and WebGL [33]. Due to use of these libraries, the Google Chrome browser is recommended to achieve the full DPLab functionality. When users upload new images, the server will automatically convert the original uploaded data files into Deep Zoom Image (DZI) format compatible with the OpenSeadragon JavaScript library used for browser-based visualization [29]. The DZI format is advantageous for web browser access as it minimizes the memory consumption needed for large image visualization by pre-tiling each original image into a pyramidal scheme of sub-images. As users zoom, pan, and navigate across an image in their web browsers, the JavaScript library running within the browsers dynamically refreshes and retrieves subsets of image tiles appropriate for the users’ current field of view and level of zoom.

The image analysis functionality of the application uses Redis and Sidekiq (i.e., a Ruby library) to manage the analysis job queue and perform tasks in a robust and asynchronous manner [49, 50]. When a user queues an analysis, a job object containing analysis options, parameters, and a pointer to the stored image file is placed into a simple Redis key-value list which functions as a job queue. A remote computational server, separate from the web server, maintains a pool of Sidekiq worker threads that continuously pulls jobs from the Redis job queue and processes them accordingly. The extensible open library of algorithms supports a variety of publicly contributed algorithms in various programming languages, including Java, MATLAB, Python, C, C++, Julia, among others. Depending on the algorithm queued for analysis, which is set as configuration, the workers call into scripts and functions based on that configuration and convert the function output into a unified format that can be saved to the database and visualized by the web application. Figure 5 outlines the overall DPLab architecture and its affiliated services and servers as discussed.

**Figure 5.**
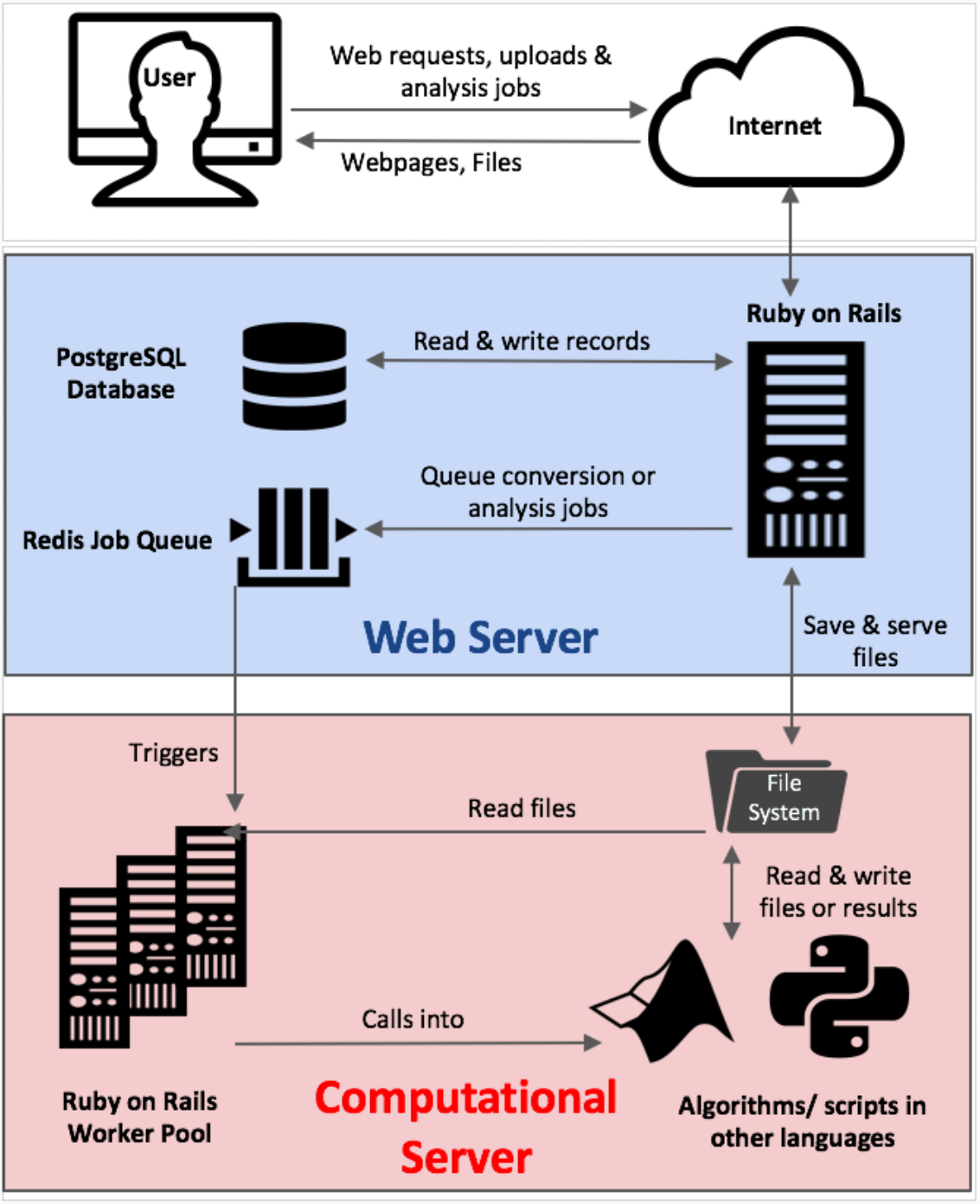
Digital Pathology server architecture and design. The splitting architecture of the DPLab server separates the web server (blue) from the computational server/cluster (red). A unified job queue and database hosted on the web server is used to manage a clustered pool of worker threads on the computational server for analyses in parallel. The architectural design of separating the computational server from the application server preserves the web application performance and provides consistent user experience, even under high computational load. It also allows for flexible scaling of the computational cluster by managing computational worker threads asynchronously, allowing new machines to be easily added to or removed from the cluster.

The architecture of the DPLab platform is designed to be highly scalable to handle the expected server load from running computationally expensive image analysis jobs by multiple users simultaneously in parallel. The bifurcation architectural design also allows DPLab to be deployed in a single optimized machine or container. There are several critical architectural features designed to improve the scalability and stability of the platform even under high computational load, as demonstrated in Figure 6. First, the computational server and the web server are architected to function on separate machines. This bifurcation prevents computational utilization of the analysis jobs from interfering with the performance of the user-facing web application. Second, the computational server uses a “worker pool” design pattern which allows efficient and parallelizable performance of analysis tasks under high load. This architecture implements a single, light-weight job queue to hold invocation data for analysis tasks originating from any number of clients, as well as a pool of worker machines to perform such jobs. A worker machine is defined as any external machine that is authorized to listen to the job queue as a worker node/server often contained within the same internal or virtual private network. The number of machines within this authorized worker pool can be readily and dynamically changed in response to fluctuating task load. Thus, this design pattern permits any number of remote machines or virtual environments with access permissions to “listen” to a synchronized job queue and act as a worker machine in performing multi-threaded computational tasks and returning results to a database cluster. Under high load conditions, the database itself can be clustered across many individual databases running on separate servers to allow for increased numbers of concurrent connections to worker threads. Using PostgreSQL’s synchronous data replication functionality and cluster support, the individual databases can consistently keep in-synchronized with other database members of the cluster by performing replication checks on a frequent and pre-defined interval. This architectural feature allows for any number of machines and threads to scale up processing of computational tasks in parallel. As server load fluctuates with demand, multiple computational analysis-performing machines, as well as database servers can be flexibly clustered together to extend and retract server performance dynamically as appropriate to “elastically” respond to such demand.

**Figure 6.**
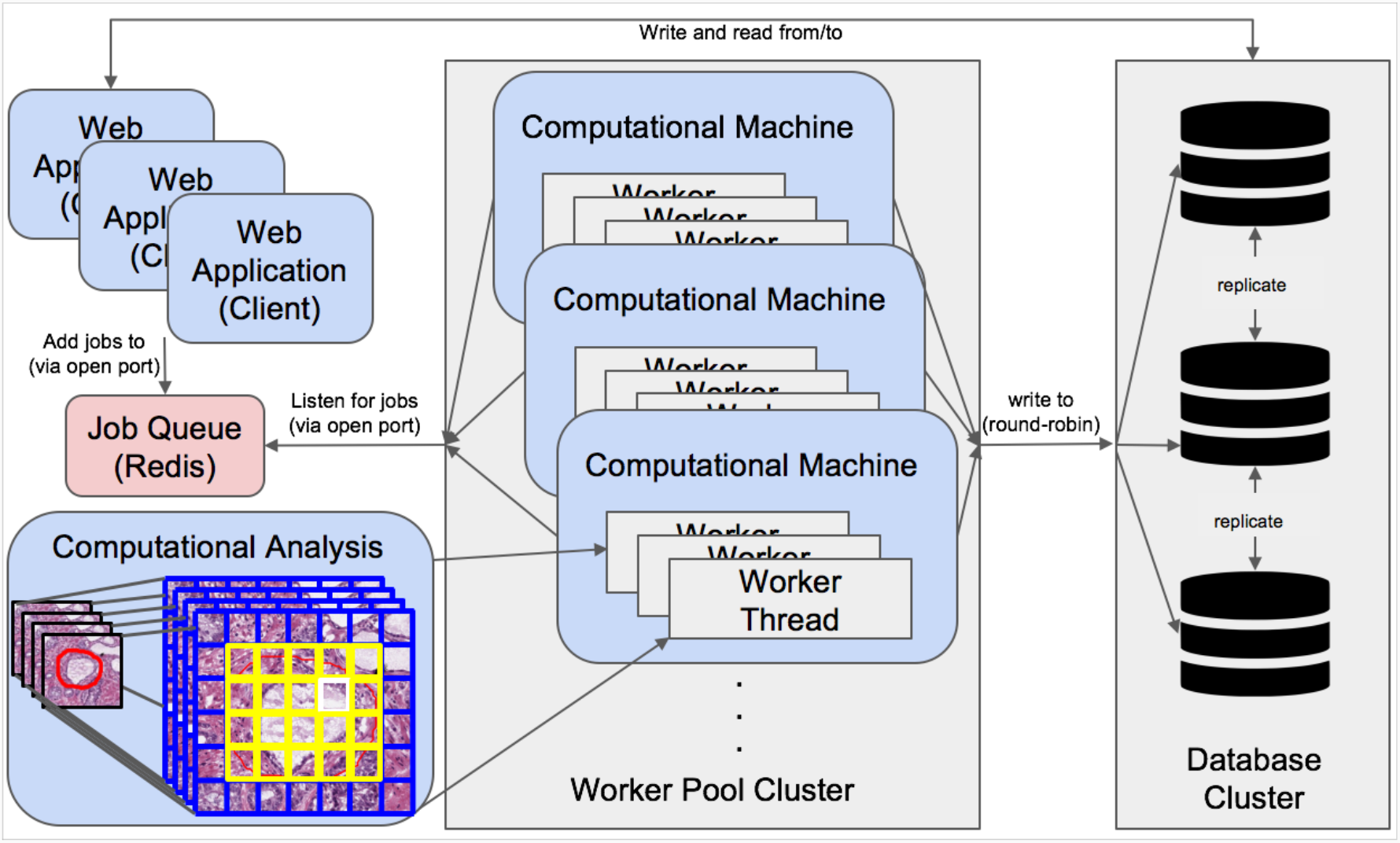
Scalable worker and database architecture. A single unified Redis job queue is used to hold and manage jobs in a first-in-first-out priority queue on the primary web application server. A scalable pool of any number of remote worker machines with access permissions to listen to the job queue on a specific network port can retrieve jobs from the application server Redis queue and perform them asynchronously in parallel. Computed data is next written to a separately hosted cluster of database servers. Database connections from computational threads are round-robined across database servers, and databases are kept in-synchronized with Postgres built-in master-slave replication functionality. The number of machines within the database cluster and computational worker pool is scalable and flexible to real-time demand.

## CONCLUSION

The developed DPLab web application is significant to tissue based cancer and pathology research community as it enables 3D histopathology image analysis, and introduces a suite of bioinformatics tools for 2D histopathology image processing. DPLab serves as an online workbench to facilitate the end-to-end analysis pipeline of pathology image data in a variety of formats and modalities, including pathology WSIs and Imaging Mass Cytometry images. In particular, it supports the cutting-edge large-scale serial whole-slide microscopy image registration, pioneers the effort on 3D tissue volume reconstruction for 3D microscopy image analysis, and serves as a new vehicle for explorations of clinically relevant 3D histopathology features and spatial topology signatures from the information-lossless 3D tissue space. Using DPLab, users can upload and store their image data in a cloud-based server, remotely run computationally-expensive analysis algorithms using a distributed server cluster and export and share their results for further use. Users can either upload their algorithms or use DPLab analysis algorithm library as a code-free, and “wrapper” web interface, thereby encouraging usage, distribution, and validation of new algorithms in both research and clinical contexts. The platform addresses critical needs in the field of digital pathology by providing a publicly-accessible data analysis and management web service to the research and clinical community at large. We intend to continue the development of DPLab by providing more analysis integration features across imaging, clinical, and genomic data, expanding our library of available open-source algorithms, and improving support for more programming languages and image formats.

## ACKNOWLEDGEMENT

This research was supported by grants from National Institutes of Health U01CA242936, R01EY028450, R01CA214928, R01CA247905, National Science Foundation ACI 1443054 and IIS 1350885, and CNPq.

## AUTHOR CONTRIBUTIONS

A.S., F.W. and J.K. conceived the original idea; A.S. and J.K. designed the research; A.S., S.P., D.B., J.A. and J.K. performed the research; All authors were involved in planning for the project; A.S. and J.K. worked on the manuscript with support from S.P., F.W, A.B.F, G.T., and D.J.B. All authors reviewed the final manuscript. A.S. and J.A. were involved in this work during internship.

## CONFLICT OF INTERESTS

The authors declare no conflict of interests.

## CODE AND DATA AVAILABLILITY

Codes are available at GitHub repository: https://github.com/jkonglab/DigitalPathology. Installation instructions are provided in detail. In addition to the presentations of the key aspects of DPLab web application in this manuscript, we provide supplementary videos for DPLab platform demonstration in details by the following link: https://jkonglab.github.io/DigitalPathology.

## COST AND MAINTENNACE

The DPLab application is currently deployed under Georgia State University domain, which is being handled by Georgia State University IT team. The application is maintained by an in-house software development team. Being an open-source tool, DPLab application welcomes community contribution and will remain cost free.

## ADDITIONAL INFORMATION

We provide three different ways for users to experience DPLab.

1. We have provided a live demonstration website for the 3D liver fibrosis analysis with results viewed in a 3D virtual tissue space: https://dplab.gsu.edu/demo. This live demonstration does not require any login credential.
2. In addition to the presentations of the key aspects of DPLab web application in this manuscript, we provide supplementary videos for in-depth DPLab platform demonstrations by the following link: https://jkonglab.github.io/DigitalPathology/. On this webpage, we present the following videos for demonstration.

Movie S1: User Account Creation

This video demonstrates how to create an account on the DPLab web application that prevents automated malicious threats for enhanced data security.

Movie S2: Project Management

This video demonstrates how to create projects and set properties of projects for different research investigations.

Movie S3: Image Management

This video demonstrates how to upload, combine, and search images in projects.

Movie S4: Image Annotations

This video demonstrates how to use annotation tools for annotations on DPLab web platform.

Movie S5: Annotation Files

This video demonstrates the resulting annotation Jason and ImageScope compatible xml files from DPLab web platform.

Movie S6: WSI Registration Analysis

This video demonstrates how to invoke image registration analysis to spatially align serial WSIs and enable 3D tissue investigations.

Movie S7: 3D Tissue Volume Visualization

This video demonstrates the 3D visualization tools for pathology WSI volume inspection.

Movie S8: Algorithm Invocation

This video demonstrates the general procedure to invoke and submit analysis jobs.

Movie S9: Algorithm Deployment

This video demonstrates how to integrate and manage new algorithms to expand the analysis library on DPLab web platform.

Movie S10: Analysis Result Visualization, Refinement, and Export

This video provides a demonstration on result review, refinement, and export.

Movie S11: Administrator Account

This video demonstrates how to manage user accounts with an administrator account.

## Notes

### Competing Interest Statement

The authors have declared no competing interest.

